# The HVCN1 channel conducts protons into the phagocytic vacuole of neutrophils to produce a physiologically alkaline pH

**DOI:** 10.1101/003616

**Authors:** Adam P. Levine, Michael R. Duchen, Anthony W. Segal

**Affiliations:** Division of Medicine, University College London; Department of Cell and Developmental Biology, University College London

## Abstract

Activation of the NADPH oxidase (NOX2) of the phagocytic vacuole of neutrophils is essential for innate immunity. Sustained activity of the oxidase requires that charge movements across the membrane are balanced. A role for the proton channel, HVCN1, has been proposed but not proven. Using the ratiometric pH indicator SNARF, introduced into the cytosol and separately into the vacuole coupled to *Candida*, we used confocal microscopy to measure changes in pH in these two compartments in human and mouse neutrophils. Shortly after phagocytosis by human cells, the vacuolar pH rose to ~9, at which it was maintained for ~20 minutes, while the cytosol showed a small acidification of ~0.25 pH unit. Alkalinisation has important consequences for the microbicidal and digestive functions of vacuolar enzymes. In HVCN1 knock out mouse neutrophils, the phagocytosis induced respiratory burst was halved to ~3 fmols per *Candida*, the vacuolar pH rose to >11 and the cytosol acidified excessively to pH ~6.75. These changes were prevented by the protonophore CCCP. The rate of extrusion of protons into the extracellular medium following phagocytosis was not significantly different from wild type neutrophils suggesting that cytoplasmic acidification resulted from the loss of the proton sink into the vacuole. HVCN1 phagocytic vacuoles showed considerable swelling, and this was blocked by CCCP and decreased by valinomycin. Stoichiometric considerations indicated that the HVCN1 channel compensates 90–95% of the oxidase-induced charge in normal cells, and in its absence, charge is carried by ions other than protons, including K^+^.

## Introduction

**N**eutrophils that encounter a bacterium or fungus engulf it into a phagocytic vacuole of invaginated plasma membrane, into which cytoplasmic granules release their contents of potentially lethal enzymes (Figure 1). These processes are associated with a burst of non-mitochondrial respiration in which electrons are passed across the membrane of the vacuole by an NADPH oxidase, NOX2, that generates superoxide [1]. This electron transport is essential for efficient killing of the microbes as evidenced by the severe immunodeficiency syndrome of Chronic Granulomatous Disease (CGD) in which the function of NOX2 is absent or compromised [2].

**Figure 1:**
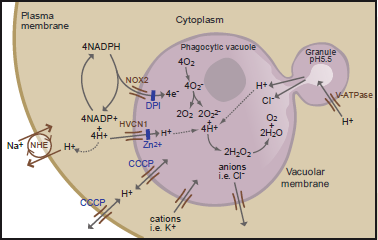
Schematic representation of the neutrophil phagocytic vacuole showing the consequences of electron transport by NOX2 onto oxygen. The proposed ion fluxes that might be required to compensate the movement of charge across the phagocytic membrane together with modulators of ion fluxes are shown.

The transport of electrons into the phagocytic vacuole is electrogenic, causing a large, rapid, membrane depolarisation which will itself curtail further electron transport unless there is compensatory ion movement [3] by the passage of cations into the vacuole and/or anions in the opposite direction (Figure 1). The nature of the ions that compensate the charge will have a direct effect on the pH within the vacuole and the cytosol. Attempts to characterise mechanisms of charge compensation have concentrated on the role of proton channels [4], characterised using divalent cations such as Zn^2^+ and Cd^2^+ as inhibitors [3], [5], [6]. These ions are reasonably selective for the proton channel in the low micromolar range [4], [7], but have multiple other targets when used at millimolar concentrations [8] – [12]. Cloning of the gene for the proton channel HVCN1 [13], [14], and the subsequent generation of HVCN1 knockout mice [15] has allowed a more precise definition of its role in neutrophil biology. Contrary to predictions from previous studies using high concentrations of Zn^2^+ [8], complete eradication of the HVCN1 channel only reduced oxidase activity by about 50% [15], [16], and had a surprisingly small effect on microbial killing [15].

Inhibition or deletion of HVCN1 channels has been shown to result in exaggerated acidification of the cytosol after phagocytosis of zymosan [17] or stimulation of the oxidase with phorbol myristate acetate (PMA) [18] which led to the suggestion that this channel might be important for the expulsion of protons from neutrophils [17], [18], although this was not measured directly in either of these studies. Those observations raise the possibility that the depressant effect of the loss of the HVCN1 channel on the NADPH oxidase might be due to the development of an excessively acidic cytosol, which is known to inhibit it [19], rather than as a consequence of impaired charge compensation.

In a subsequent study of HVCN1^-/-^ neutrophils phagocytosing zymosan particles [20] it was observed that when the pH of the of the entire population of phagocytic vacuoles of wild type and HVCN1^-/-^ neutrophils were compared, no significant difference was observed. Minor differences in the proportions of alkaline, neutral and acidic vacuoles by genotype were seen.

In order to define the role of the HVCN1 channel in neutrophil biology, we have used the ratiometric fluorescent pH indicator, SNARF, to measure the pH directly in the phagocytic vacuole and in the cytosol of human, and wild type (WT) and HVCN1^-/-^ mouse neutrophils during phagocytosis. The data confirm that the HVCN1 channel plays a major role in charge compensation across the phagocytic vacuole membrane, and in its absence non-proton fluxes predominate in charge compensation, but do not support its proposed importance for pH regulation across the plasma membrane.

## Results

### The respiratory burst is impaired in HVCN1^-/-^ neutrophils whilst extracellular acid release is normal

In WT neutrophils, oxygen consumption was increased 10 fold by PMA and to a similar level by the addition of opsonised *Candida*. PMA stimulated oxygen consumption was reduced to 68% of normal in cells isolated from HVCN1^-/-^ mice, consistent with previous observations [15], [16], was abolished by the oxidase inhibitor DPI, and was absent in neutrophils lacking gp91^phox^ (Figure 2A). Oxygen consumption by HVCN1^-/-^ cells phagocytosing opsonised *Candida* was 50% of that by wild type neutrophils, and significantly lower than in the same cells after PMA stimulation (p = 0.003).

**Figure 2:**
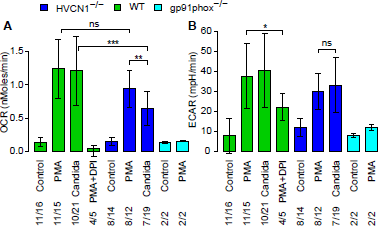
Oxygen consumption rate (OCR) (A) and extracellular acidification rate (ECAR) (B) by neutrophils from WT, HVCN1^-/-^ and gp91^phox-/-^mice in response to stimulation with PMA (with and without DPI) or opsonised *Candida*. The numbers of independent experiments is shown over the total number of measurements. Statistical significance: *** p<0.001, ** p<0.01 and * p<0.05. Differences between PMA stimulated WT and gp91^phox-/-^ were p<0.001 in (A) and p<0.05 in (B). There were no significant differences in (B) between PMA and *Candida* or between WT and HVCN1^-/-^ cells. Mean ± SD shown.

Activation of the oxidase using PMA or *Candida* increased the extracellular acidification rate (ECAR), to similar levels in HVCN1^-/-^ and WT mouse neutrophils and levels achieved were equivalent with PMA or with *Candida* (p = ns for all comparisons). The rate was reduced in the presence of DPI and remained at baseline levels in neutrophils lacking gp91^phox^ (Figure 2B).

Oxygen consumption by human neutrophils phagocytosing *Candida*, as measured with a Clark oxygen electrode, was determined as 5.8 fmols ± 0.4 (SD) (n=5) per *Candida* phagocytosed and the respiratory burst was brief, with a peak rate lasting about two minutes [21]. In HVCN1^-/-^ the equivalent measurement was 3.2 fmols corresponding to the 50% reduction observed (Figure 2A). Given the 4:1 ratio of electrons passing across the vacuolar membrane to oxygen consumed (Figure 1), the consumption of ~6 fmols of oxygen per *Candida* would require ~24 fmols of compensating charge in normal neutrophils and ~12 fmols in HVCN1^-/-^ cells.

### The phagocytic vacuole of human and WT mouse neutrophils reaches pH 8.5-9 whilst in HVCN1^-/-^ it exceeds 11

Shortly after an organism was engulfed, the vacuole underwent a significant alkalinisation (Figure 3). In human neutrophils, this started almost immediately (Figure 3E and Video S1). The mean maximum pH reached post-phagocytosis was 9 (Figure 3E) and in most cells this elevated pH was maintained for 20-30 minutes. When the NADPH oxidase was inhibited by DPI, the vacuole acidified from the outset (Figures 3B, 3E) and when added following phagocytosis it rapidly reduced the vacuolar pH (Figure 4C and Video S2) [20].

**Figure 3:**
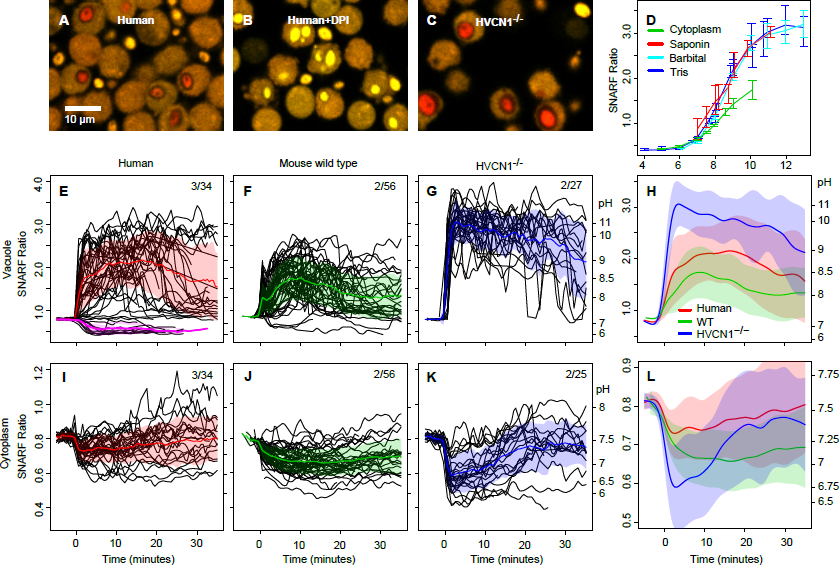
Time courses of changes in pH in the vacuole and cytoplasm of phagocytosing neutrophils. Representative images of *Candida* phagocytosed by human neutrophils, without (A), and with DPI (B), and by HVCN1^-/-^ neutrophils (C). Standard curves for the relationship between SNARF ratio and pH of organisms and cytoplasm are shown in (D). *Candida* alone were added to two different buffer systems, labelled Tris or Barbital, and intracellular organisms were exposed to the Barbital buffers after permeabilisation of neutrophils with saponin. Panels E-L show time courses of the pH changes of phagocytosed *Candida* and cytoplasm of human (E, I), mouse WT (F, J) and HVCN1^-/-^ (G, K) neutrophils synchronised to the time of particle uptake (0 minutes). In E-G and I-K, each individual black line represents serial measurements of the SNARF ratio of a single phagocytosed *Candida* or neutrophil cytoplasm, respectively. Mean ± SD (shaded areas) are shown. In the composite panels H and L the mean data have been smoothed. Data are plotted according to SNARF ratio with the approximate corresponding pH shown on the right y-axis. The number of independent experiments over the total number of individual cells examined is shown. The effect of DPI is shown in E (pink, 11 cells).

**Figure 4:**
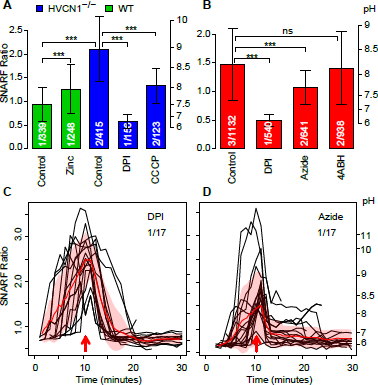
The effect of DPI, azide, 4ABH, zinc, and CCCP on vacuolar pH. (A) The effect of 100 µM Zn^2^+, DPI and 60 µM CCCP on vacuolar pH in WT and HVCN1^-/-^ neutrophils at ~30 minutes following the addition of *Candida* without synchronisation to particle uptake. (B) The effect of 5 µM DPI, 5 mM sodium azide and 50 µM 4ABH on vacuolar pH in human neutrophils at ~30 minutes following the addition of *Candida* without synchronisation to particle uptake. The effect of the addition of azide (C) or DPI (D) on vacuolar pH in human neutrophils synchronised to time of addition (red arrows).The numbers of experiments is shown over the number of independent measurements. Mean ± SD shown. Statistical significance: * * * p<0.001.

These elevations in vacuolar pH occurred despite the buffering capacity of the heat-killed *Candida* (Figure S1). The maximum difference in the vacuolar pH in human neutrophils in the presence or absence of DPI (~6.5 and ~9, respectively (Figure 3E)) required 1.9 fmols of OH^−^ per *Candida*. Assuming this pH rise was entirely due to nonproton charge compensation, it would amount to ~8% of the total ~24 fmol compensating charge, close to the ~5% previously estimated [22].

WT mouse neutrophils showed a similar, although less extreme, alkalinisation of the vacuole reaching a mean maximum pH of ~8.5 post-phagocytosis (Figure 3F). The initial rise was very rapid and was followed by a brief fall, possibly as a result of degranulation of acid granule contents, before resuming its upward course after a delay of about one minute. By contrast, the pH in the vacuoles of HVCN1^-/-^ neutrophils was rapidly and grossly elevated to a mean value of ~11.1 (Figure 3G and Video S3). These changes were observed in almost all vacuoles. Given the strong buffering capacity of *Candida* in the alkaline range, a rise in pH from ~6.5 in DPI treated cells to ~11.1 in HVCN1^-/-^ required ~9.2 fmols of OH- per organism (Figure S1), similar to the ~12 fmols of total compensating charge.

In a small number of HVCN1^-/-^ cells the vacuolar membrane was unable to contain the alkalinity, causing the vacuole to collapse (e.g. Videos S3 and S4), which coincided with sudden alkalinisation of the cytoplasm. These extreme conditions could impair the oxidase, accounting for the lower rates of oxygen consumption produced by phagocytosing HVCN1^-/-^ neutrophils.

The alkalinisation in HVCN1^-/-^ vacuoles was almost completely prevented by the addition of the protonophore carbonyl cyanide-m-chlorophenylhydrazone (CCCP), whereas Zn^2^+, increased alkalinisation of vacuoles in WT mouse neutrophils at a concentration of 100 µM (Figure 4A).

Sodium azide has been used [20], [23] to inhibit the action of myeloperoxidase (MPO) which was thought to quench the fluorescence of fluorescein, the flurophore previously utilised to determine vacuolar pH. We therefore determined the effect of the addition of azide on SNARF fluorescence. Azide reduced the pH in the vacuoles at ~30 minutes in human neutrophils (Figure 4B) or almost immediately when added during a time course (Figure 4D). This effect of azide was not due to its inhibitory effect on MPO because 4-aminobenzoic acid hydrazide (4ABH), a selective inhibitor of MPO, had no effect on vacuolar pH at 50 µM, more than three times the IC50 [24] (Figure 4B). When SNARF-labelled *Candida*, were added to buffers at varying pH in the absence or presence of azide, no effect was seen on the SNARF ratio (Figure S3).

### Cytoplasmic pH falls after phagocytosis, and this is exaggerated in HVCN1^-/-^ cells

In human or WT mouse neutrophils the respiratory burst was associated with a modest acidification of the cytoplasm, from a pH of ~7.5 to a mean nadir pH of 7.3 (Figure 3I and Video S1), with homeostasis bseing achieved over the ensuing 20–30 minutes as described previously [25] – [27]. Neutrophils from WT mice behaved similarly (Figure 3J). In HVCN1^-/-^ neutrophils the acidification was much more pronounced (p = 2x10^−5^, Figure 3K and Video S3), reaching a minimum mean pH of ~6.7 at 2.5 minutes after phagocytosis, before returning to normal after about 20-30 minutes. The rate of recovery of cytoplasmic pH after acidification with 20 mM NH_4_Cl was no different between WT and HVCN1^-/-^ neutrophils (Figure S4).

The buffering capacity of the neutrophil cytosol has previously been determined at between 28 [28] and 50 [26] mM unit^-1^. Given an approximate neutrophil volume of ~300 μm^3^ [29] and the 0.2 unit fall in cytoplasmic pH in human neutrophils (from 7.5 to 7.3) this equates to the accumulation of ~2.3 fmols (39 mM unit^-1^ x 0.2 unit x 300 μm^3^) of protons left in the cytosol when the charge is compensated by alternative ions (9.6% of ~24 fmol). In HVCN1^-/-^ cells the fall in cytoplasmic pH to 6.6 equates to ~10.5 fmol of protons (calculated as above).

### HVCN1^-/-^ vacuoles are larger than

The vacuoles of HVCN1^-/-^ neutrophils containing *Candida* underwent a profound swelling, which was much more obvious than in WT mouse or human cells (Videos S1 and S3). To determine the extent of this swelling and to distinguish it from that induced by the osmotic effects of the products of digestion of the *Candida*, we measured the cross-sectional areas of vacuoles containing a single indigestible latex particle with a diameter of 3 μm (7.1 µm^2^), similar to that of *Candida*. In neutrophils from HVCN1^-/-^ mice, the vacuoles swelled to a mean cross-sectional area of 13 μm^2^ as compared with 8.2 μm^2^ in neutrophils from WT mice (p<10^−26^) or HVCN1^-/-^ treated with 5 μM DPI (7.7 µm^2^) or CCCP (7.9 µm^2^) (Figure 5) after 30 minutes. These results indicate that osmotically active ions are driven into the vacuole by the oxidase in the absence of the proton channel. This swelling was reduced by 3 μM valinomycin to a mean area of 11 µm^2^ (p<10^-5^). The estimated volumes utilising the radius calculated from the median cross-sectional area and assuming spherical vacuoles were: latex particles (14.2 µm^3^); WT vacuoles (17.6 µm^3^) and HVCN1^-/-^ (35.4 µm^3^) without and (27.4 µm^3^) with valinomycin, respectively.

**Figure 5:**
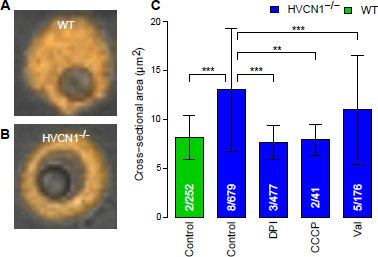
Vacuolar size in WT and HVCN1^-/-^ neutrophils containing a single latex (cross-sectional area ~7 µm^2^) particle. Representative images are shown in (A) and (B). (C) Quantitation of vacuolar swelling in HVCN1^-/-^ neutrophils compared with WT, and the effects of 5 µM DPI, 60 µM CCCP and 3 µM valinomycin. The numbers of independent experiments is shown over the total number of measurements. Mean ± SD shown. Statistical significance: *** p < 0.001 and ** p<0.01.

If the charge compensation in HVCN1^-/-^ cells (12 fmols) were purely accomplished by the influx of cations, e.g. K+, to a concentration that balanced the cytosolic osmotic pressure of 230–235 mOsm [30] it would produce a vacuolar volume of ~52 µm^3^ (12 fmols / 232.5 mM) considerably larger than that observed.

## Discussion

The original description [31] of an alkalinisation of the neutrophil phagocytic vacuole by the NADPH oxidase to a pH of ~7.5-8 has been replicated a number of times [32] – [35]. In those studies, fluorescein was employed as pH indicator. Fluorescein saturates at a pH of ∼8 and above, was not generally used in a ratiometric manner. In addition, it has been thought by some to become bleached by the action of myeloperoxidase in the phagocytic vacuole [36], [37].

SNARF is much better suited to the determination of the vacuolar pH. It has a dynamic range of between 6 and 11 (Figure 3D), is ratiometric, and can also be used to simultaneously measure the pH in the vacuole and cytoplasm. We have shown in several ways that its fluorescent properties are not altered by the conditions in the phagocytic vacuole. The extent and duration of alkalinisation we observed in the human neutrophils was much greater than that previously described by us [31] and others, because SNARF is more suited to measurements in the alkaline range. In addition, we only measured the pH of *Candida* inside vacuoles, so that the fluorescence signal was not influenced by that of the non-phagocytosed organisms that are buffered by the extracellular medium. Synchronising the changes in fluorescence to the time of particle uptake, gave a more accurate indication of the changes in pH over time.

Both studies that were unable to detect an elevation of neutrophil vacuolar pH above neutral [20], [23] included azide in all the solutions to counteract bleaching of the fluorescein by MPO [23]. We found here that azide significantly reduced the vacuolar pH (Figure 4B and 4D) by a mechanism that did not involve the inhibition of MPO; 4ABH, which is a more specific MPO inhibitor, had no such effect at a concentration of over three times the IC50 [24] and the addition of azide produced a rapid drop in the pH of previously alkaline vacuoles implying a direct effect of azide on pH (Figure 4D). Sodium azide has known proton uncoupling capabilities [38], [39] and might depress vacuolar pH by acting as a protonophore across the vacuolar membrane in much the same way as does CCCP (see below).

The prolonged phase of alkalinisation of the phagocytic vacuole to between 8.5 and 9.5 would provide an optimal milieu for the microbicidal and digestive functions of the major granule proteases, elastase, cathepsin G and Proteinase 3 [22], [40]. The peroxidatic action of MPO has a pH optimum of ~5 and is minimal above neutral so the possibility of the efficient generation of HOCl at pH 9 is low [41], [42].

The gross difference in the vacuolar pH between HVCN1^-/-^ neutrophils and those from WT mice and humans must indicate that less protons pass into the vacuole in the HVCN1^-/-^ cells, providing firm evidence that under physiological conditions the HVCN1 channel compensates the oxidase induced charge by conducting protons across the vacuolar membrane. The excessive acidity that develops in the cytoplasm in HVCN1^-/-^ neutrophils, despite a normal proton extrusion rate out of the cell, is further evidence that the protons generated by activity of the oxidase on the cytosolic side of the vacuolar membrane accumulate because they do not pass into the vacuole. Additional proof that it is the failure of the movement of protons into the vacuole that causes the extreme alkalinisation in the HVCN1 phagocytic vacuoles is provided by the effect of the addition of the protonophore, CCCP, which overcomes the barrier to proton flux and largely reverses the alkalinisation. These results provide direct evidence for compensation by HVCN1 channel of the NADPH oxidase induced charge across the vacuolar membrane.

Two studies have concluded that the HVCN1 channel plays an important role in controlling the cytoplasmic pH by extruding protons into the extracellular medium. The first [18] used 2,7-bis-(carboxyethyl)-5-(and-6)-carboxyfluorescein (BCECF) as indicator in HVCN1^-/-^ cells which had been suspended in a sodium free medium to prevent the activity of Na+/H+ exchangers [43], [44] which are known to regulate cytoplasmic pH. The cells were stimulated with PMA, which leads to activation of the oxidase at the plasma membrane, where both charge compensation and restitution of the pH should occur. The second study [17] measured cytosolic pH with SNARF in human neutrophils phagocytosing zymosan in the presence of 100 µM Zn^2^+. In these experiments it was uncertain what effect the Zn^2^+, and/or the 10% ethanol in which the cells were bathed for 15 minutes, might be having on cellular processes other than proton transport through the HVCN1 channel. HVCN1^-/-^ cells recovered from frozen bone marrow were also studied, but the recovered ‘phagocytes’ would have been predominantly macrophages as neutrophils are very vulnerable to freezing [45].

In neither of those studies was the rate of excretion of protons into the extracellular medium measured. In the present study we have shown that proton extrusion rates by HVCN1^-/-^ cells are normal. With proton extrusion as the rate limiting step [46] the cytoplasmic pH, will be directly dependent upon the rate at which protons are generated. Consequently, in HVCN1^-/-^ cells the cytoplasm acidifies excessively because the majority of protons generated by activation of the oxidase do not move into the vacuole.

There was surprisingly good agreement between the observed oxygen consumption and pH changes and the calculations of the stoichiometry of the required compensating charges and the proportions of these contributed to by protons, 5-10% in normal cells and >90% in HVCN1^-/-^. However, the vacuolar swelling observed was less than would be expected were all the non-proton mediated charge compensation to be due to the influx of K+, indicating that part of the charge compensation in these cells may be due to the efflux of anions, e.g. Cl^−^, from the vacuole. A role for K+ influx [22] is supported by our observations of the reduction in vacuole swelling by valinomycin and previous data on Rb+ efflux from PMA stimulated neutrophils [47], [48].

Cytoplasmic granule acidification requires the vacuolar-type H+-ATPase to pump protons into the lumen of the granules [49] and a counter-ion flux would be required to neutralize the membrane potential created by proton accumulation. Cl^−^ is the likely counter ion [50], which could enter the granules through CLIC1 [51]. The Cl^−^ in the granules would not provide sufficient non-proton charge compensation by itself. A granule volume of about 10% of that of the neutrophil, with a Cl^−^ concentration of ~80 mM [52], and the degranulation of a quarter of the granules into each vacuole, would only produce 1 fmol, or 10% of the required compensating charge in HVCN1^-/-^ neutrophils, so some mechanism would be required to regenerate Cl^−^ in the vacuole. Chloride movement into phagocytic vacuoles has been proposed to occur through CFTR [53] and CLCN3 [54], but this would require movement against a strong electrical gradient. KCC3 has been shown to be required for normal oxidase activity [47] and being an electroneutral co-transporter of K+ and Cl^−^, it could satisfy the dual requirements of moving chloride into the vacuole to be cycled out through a chloride channel whilst leaving K+ in the vacuole to activate the granule enzymes [22].

## Materials and Methods

### Materials

SNARF-1 was from Invitrogen. Other chemicals, and latex particles, were from Sigma

### Media

Balanced salt solution (BSS) contained 156 mM NaCl, 3.0 mM KCl, 1.25mM KH_2_PO^4^, 2 mM MgSO_4_, 2 mM CaCl_2_, 10 mM glucose, 10mM Hepes at pH 7.4. The KH_2_PO^4^ was replaced with KCl in experiments employing Zn^2^+. Phosphate buffered saline (PBS) was from Gibco.

### Organisms

*Candida, albicans* was a clinical isolate.

### Labelling *Candida* with SNARF

*Candida* were washed twice, resuspended in PBS, heated to 60° C for 30 minutes and washed and resuspended at 1x10^8^/ml in 0.1 M NaHCO_3_ pH 8.5. 50 µg SNARF-1 succinimdyl ester in 100 µl dimethyl sulphoxide (DMSO) was added drop wise to 2 ml of a rapidly stirred suspension of these cells at room temperature. After mixing for 30 minutes at room temperature, the labelled cells were washed twice, and then resuspended in 2 ml of BSS.

### Particle opsonisation

C57B6 mice were injected three times over 6 weeks with 1x10^7^ heat killed (60°C, 30 minutes) C. albicans. 1x10^7^ SNARF labelled *Candida*, in 100μl PBS were incubated with 10% complement preserved serum (Patricell) and 50 μl immune serum for 60 minutes at 37° C. Latex particles 3 μm (Sigma) in 0.1 M NaHCO3 pH 8.5 were opsonised with an equal volume of normal mouse IgG (Caltag) overnight at 4°C. For studies on human neutrophils, *Candida* were opsonised as above with 50% human IgG (160 mg/ml Vivaglobulin CSL Behring) plus 10% pooled normal human serum. All particles were washed and resuspended in 1 ml BSS.

### Isolation of neutrophils

Thioglycollate-elicited peritoneal neutrophils were collected from mice by peritoneal lavage 18 hours after intraperitoneal injection of 3% thioglycollate, and maintained in PBS with 5 U/ml heparin. The cells were centrifuged through Lymphoprep (density 1.077, Axis-Shield) at 400 g for 10 minutes and then washed and resuspended in BSS. Peripheral blood was obtained from mice by cardiac puncture, taken into heparin (5 U/ml) and immediately diluted with an equal volume of PBS with heparin (5 U/ml). This mixture was then diluted tenfold with PBS and one tenth volume of 10% dextran was added. After sedimentation for one hour the supernatant was removed, centrifuged through Lymphoprep (density 1.077) at 400 g for 10 minutes. The pellet was subjected to hypotonic lysis and the neutrophils pelleted at 200 g for 10 minutes. Cells were then resuspended in BSS. All experiments were performed on pooled blood from at least two mice. Human neutrophils were purified from peripheral blood by the standard procedures of dextran sedimentation, centrifugation through Lymphoprep and hypotonic lysis.

### Oxygen consumption

For the Seahorse apparatus (Seahorse Bioscience), 4×10^5^ peritoneal neutrophils in 100 µl BSS medium were added to the poly-L-lysine coated wells of 24 well microplates and incubated at room temperature for 30 minutes. 4x10^7^1 opsonised, heat-killed *Candida* or 5 µM PMA were added to each well and oxygen consumption and extracellular pH were measured repeatedly using the Seahorse XFe96 Extracellular Flux Analyzer over 15 minutes. 5 µM DPI was added to some wells immediately before the PMA. 1x10^7^ neutrophils were rapidly stirred in a Clarke type oxygen electrode (Rank Brothers) at 37°C. 1x10^8^ opsonised *Candida* were added and phagocytosis assessed microscopically.

### Confocal microscopy

Neutrophils (1-5x10^5^ in 300 µl BSS) were incubated for 30 minutes on 25 mm poly-L-lysine coated coverslips in Leiden dishes, or poly-L-lysine coated eight well ibi treated μ-Slides (Ibidi, Germany). For labelling of cytosol with SNARF, 1% by volume of 50 µg carboxy SNARF-1, AM ester, acetate in 100 µl DMSO was added for five minutes at room temperature. The cells were washed and resuspended in 400 μl BSS for the coverslip chamber and 200 μl for μ-Slides. For time course experiments, coverslips in Leiden chambers were incubated at 37° C on a heated stage and 2×10^6^ opsonised SNARF labelled organisms added. Images were taken every minute for 30-60 minutes. For end-point measurements in slide chambers the labelled cells were incubated for 15 minutes at 37° C and then 1×10^6^ particles were added and images obtained after a further 15-30 minutes at 37° C. All confocal experiments were performed on peripheral blood human or mouse neutrophils.

Cells were imaged with a Zeiss 700 confocal microscope using a 63× oil immersion. SNARF-1 fluorescence was excited at 555 nm and the emission was split between two detectors measuring fluorescence emission simultaneously between 560-600 nm and >610 nm. Only cells containing a single *Candida* or latex particle were analysed.

### SNARF calibration

Cytosolic pH standard curves were constructed by the nigericin/high K+ technique [17]. Cells were labelled with SNARF as described above and then washed and incubated in high K+ solutions (100 mM K+) containing 10 μM nigericin and 50 mM buffers at pH 3 (glycine), 4-6 (acetate), 7-9 (Tris), 10 (glycine) and 11-13 (PO_4_) in 100 mM KCl. Standard curves were also constructed with 100 mM KCl and a 50 mM mixture of barbital, citric acid, boric acid and PO4 adjusted to a pH of between 3 and 13. The cytoplasmic pH was allowed to equilibrate with each external solution until the ratio of SNARF no longer changed with time. SNARF labelled *Candida* were equilibrated in the same buffers without nigericin. Human neutrophils that had phagocytosed SNARF-labelled *Candida* were suspended in the buffers in the presence of 0.3% saponin before imaging the *Candida*.

### SNARF-labelled *Candida* provide a suitable pH indicator system for measuring vacuolar pH

Standard curves in the two buffering systems demonstrated that the dynamic range of SNARF ranged between a pH of 6 and 11 (Figure 3D). The standard curve on intracellular organisms in permeabilised human neutrophils (Figure 3D) was not significantly different. There was considerable variation of the fluorescence ratio of the individual *Candida* organisms, particularly at higher pH. The fluorescent properties of the *Candida* were not permanently altered by the conditions in the vacuole. In some cases in HVCN1^-/-^ neutrophils (Videos S3 and S4) the vacuoles ruptured, releasing the organisms, the fluorescence of which reverted to levels comparable to those of other extracellular labelled organisms. Finally, there was no decrease in fluorescence intensity at the two emission wavelengths measured over the time course of the experiments (Figure S2).

### Quantitation and statistical analyses

SNARF ratio (>610 nm / 560-600 nm) and vacuolar cross-sectional areas were determined by manual quantitation under blinded conditions using custom macro scripts in ImageJ (National Institutes of Health, USA, version >1.46r). Data were analysed using the R statistical package. For time courses, data were synchronised to the time of particle uptake and were normalised to the mean baseline pre-phagocytic ratio. Linear interpolation was performed between measurements. Statistical comparisons were undertaken by analysis of covariance (using the *lm* function) with experiment identifier as a covariate to correct for interexperiment variation.

### Mice

The knock-out mouse strains used in this study were: HVCN1 [17], and X-CGD mice B6.129S6-Cybbtm1Din/J deficient in gp91^phox^ (The Jackson Laboratory). Mice were genotyped according to previously described protocols.

## Acknowledgments

We thank the Wellcome Trust and the Irwin Joffe Memorial Fellowship for support, Phillipe Behe for very helpful discussions, Melania Capasso for providing us with the HVCN1^-/-^ mice and Adrian Thrasher for the mice lacking gp91^phox^, and Penelope Harrison, Bernadette Pedersen, Olivia Sheppard, Sabrina Pacheco, Janne Plugge, Sam Ranasinghe and Will Kotiadis for excellent technical assistance.

## Supplementary Figures

**Figure S1:**
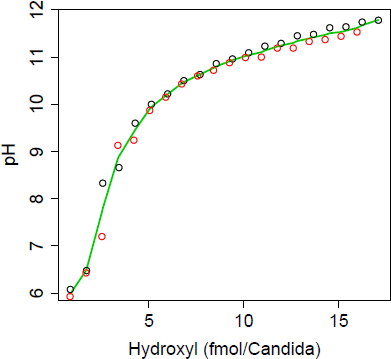
Titration of the buffering capacity of *Candida* on pH changes in response to the addition of KOH. Two independent experiments (red and black) are shown.

**Figure S2:**
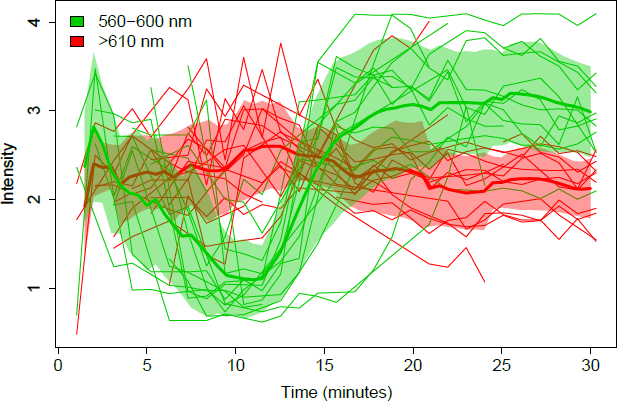
Demonstration of the time course of changes in intensity of the two emission wavelengths from the experiment shown in Figure 4B showing that there is no loss of intensity over time.

**Figure S3:**
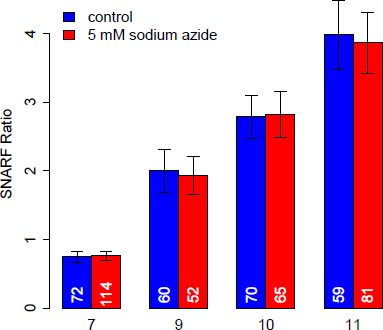
The lack of an effect of 5mM sodium azide of pH dependence of SNARF ratio. SNARF labelled *Candida* were incubated in the Barbital buffer at pHs of 7, 9, 10 and 11 for 30 min. and the fluorescence measured. The numbers of measurements are given at the bottom of each bar. Mean ± SD are shown.

**Figure S4:**
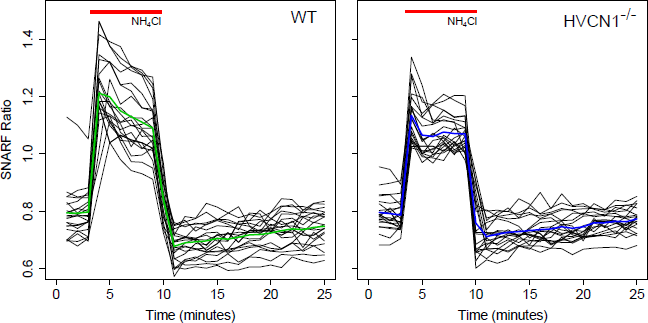
The effect of the addition of 20 mM NH_4_Cl on the ratio of SNARF in the cytoplasm of WT (left) and HVCN1^-/-^ (right) neutrophils alone. Cells were perfused with BSS. During the interval indicated by the red bars the perfusion medium was changed to a mixture of 80% BSS and 20% 0.15 M NH_4_Cl. Black lines represents serial measurements of individual cells with the mean shown in colour.

## Video Legends

**Video S1**

Time lapse video of human neutrophils phagocytosing *Candida*.

**Video S2**

Time lapse video of human neutrophils phagocytosing *Candida* with the addition of 5 μM DPI during the time course.

**Video S3**

Time lapse video of HVCN1^-/-^ neutrophils phagocytosing *Candida*. Following uptake the Candida particles turn red indicating extreme alkalinisation and the vacuoles can be seen to swell. In a small number of cells the cytoplasm suddenly becomes alkaline.

**Video S4**

Time lapse video of an HVCN1^-/-^ neutrophil phagocytosing and releasing a *Candida* particle which is subsequently phagocytosed by an adjacent neutrophil.

